# Single cell multi-omics reveals rare biosynthetic cell types in the medicinal tree *Camptotheca acuminata*

**DOI:** 10.1101/2025.04.22.650021

**Authors:** Van-Hung Bui, Joshua C. Wood, Brieanne Vaillancourt, John P. Hamilton, Lemor H. Carlton, Thu-Thuy T. Dang, C. Robin Buell, Chenxin Li

## Abstract

*Camptotheca acuminata* Decne is a woody medicinal tree that produces over a hundred bioactive compounds, including camptothecin, which has been used as the starting material to semi-synthesize many leading anticancer drugs (Lorence and Nessler 2004). Camptothecin and its derivatives are potent inhibitors of DNA topoisomerase I and are widely used for the treatment of lung, cervical, ovarian, and colon cancers. Camptothecin biosynthesis in *C. acuminata* involves complex catalytic steps, most of which remain undeciphered. In this pathway, tryptamine and secologanic acid are coupled, leading to strictosidinic acid. The formation of strictosidinic acid is catalyzed by strictosidine/strictosidine acid synthase enzymes (STR) (Fig. 1A). While a biosynthetic route for the conversion of the indole ring to the quinoline ring has been proposed, most of the underlying biosynthetic genes have yet to be identified (Fig. 1A) (Sadre et al. 2016). In addition, the cell type specificity of this pathway also remains undescribed. Here, we generated a single cell multiome (RNA-seq and Assay for Transposase Accessible Chromatin by sequencing [ATAC-seq] from the same nuclei) to probe the cell type specificity of camptothecin biosyn-thetic genes.

---

We performed organ-level metabolite profiling on key biosynthetic intermediates (Fig. S1) across multiple *C. acuminata* organs and found that camptothecin was detected across all organs tested (Fig. S1). Young leaf was chosen for single cell omics experiments (Table S1) due to the relative ease of nuclei isolation (Fig. S2A-C), which permits the simultaneous profiling of gene expression and chromatin accessibility. To aid downstream bioinformatic analyses, we produced a new version of the *C. acuminata* genome assembly and annotation (Table S2-S5, Fig. S2D), in which we used Nanopore full-length cDNAs to refine gene model predictions. For the gene expression assay of the multi-omics experiment, we obtained gene expression profiles for 4,012 high quality nuclei and 26,074 expressed genes (Table S6, Fig. S3), which includes previously characterized biosynthetic genes (Table S7) and previously reported camptothecin decorating enzymes, *CPT11H* and *CPT10-OMT* (Nguyen et al. 2021; Salim, Jones, and DellaPenna 2018).

Using unsupervised clustering and previously established leaf cell type marker genes in Arabidopsis (Table S8) (Kim et al. 2021), we identified major cell types of the leaf (i.e., mesophyll, epidermis, and vasculature) (Fig. 1B and Fig. S4A). Among the expressed biosynthetic genes (Table S7), *STR* genes were highly specific to a rare cell type that accounts for only 4.4% of all leaf cells, which we termed “STR+ cells” (Fig. 1C). Expression of key biosynthetic step(s) in a rare cell type is reminiscent of restriction of vinca alkaloid biosynthetic genes to the rare idioblast cells in *Catharanthus roseus* (Li et al. 2023). We next performed a joint RNA-ATAC analysis by matching the cell barcodes from both assays (Fig. S4B) and thus transferring the cell type identity from the RNA-seq assay to the ATAC-seq assay. The ATAC-seq signals were strongly enriched at transcriptional start and end sites (Fig. S5A) and highly enriched at ATAC-seq peaks (Fig. S5B, Table S9). Among 48,456 ATAC-seq peaks, 199 (0.41%) were detected as STR+ marker peaks (ATAC-seq peaks that were specifically enriched in STR+ cells) (Fig. 1D).

We found that among the 121 genes that were within 2-kb of a STR+ marker peak, 34 (28%) of them were most highly expressed in STR+ cells, representing 9.3-fold enrichment over the expected background ratio (Fig. 1E, *p* < 2.2 × 10^-16^, Chi-squared test). The enrichment of STR+-expressed genes suggests that STR+ marker peaks could act as cell type specific enhancers. *De novo* motif enrichment on STR+ marker peaks revealed an overrepresentation of MYB binding motifs (Fornes et al. 2019) among these accessible chromatin regions (Fig. 1F), suggesting MYB family TFs may be involved in cell type specific expression of *STR* genes in *C. acuminata*.

**Fig. 1.**
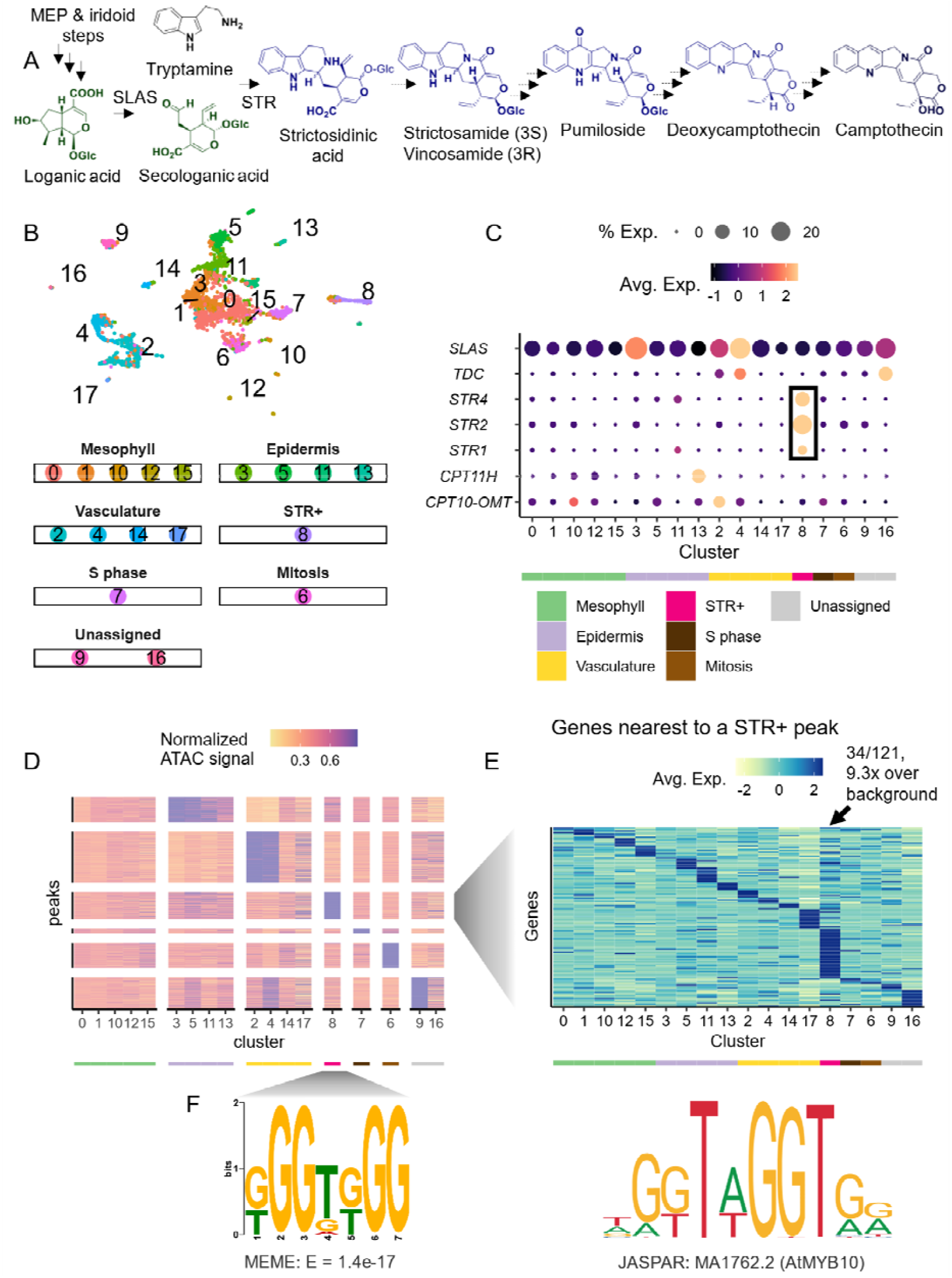
Single cell multi-ome of *Camptotheca acuminata* leaf. (A) The proposed biosynthetic pathway for camptotheci . Solid arrows indicate previously characterized enzymatic steps. Dashed arrows indicate proposed enzymatic steps (see Table S7 for gene name abbreviations). (B) Uniform Manifold Approximation and Projection (UMAP) of nuclei of the single nuclei RNA-seq dataset (n = 4,012), color coded by cell clusters. (C) Gene expression heatmap of MIA biosynthetic genes across cell clusters. Rows are expressed biosynthetic genes, which are ordered from upstream to downstream. Color scale shows the average scaled expression of each gene at each cell cluster. Cell clusters are sorted by cell types. Dot size indicates the percentage of cells where a given gene is detected. The predicted cell type for each cell cluster is annotated by the color strip below the x-axis. Box highlights expression of STR genes. (D) Heat map showing accessibility of cell type marker peaks across cell clusters. Each row is an ATAC-seq peak. Each column is a cell cluster. Color scale is maxed out at 90^th^ percentile of normalized ATAC-seq signal. The predicted cell type for each cell cluster is annotated by the color strip below the x-axis, with the same color palette as (B). (E) Heatmap showing gene expression across cell clusters. Each row is a gene within 2-kb of a STR+ marker peak. Each column is a cell cluster. The predicted cell type for each cell cluster is annotated by the color strip below the x-axis, with the same color palette as (B). (F) DNA motif enriched in STR+ marker peaks, as well as a reference MYB motif.

Taken together, our results showcased that single cell omics techniques are powerful in detecting rare biosynthetic cell types or cell populations and the regulatory landscape within. The datasets generated by this study are valuable resources for mining new biosynthetic genes and *cis*-regulatory elements for camptothecin biosynthesis, and underscore the potential of single-cell technologies in natural product research beyond herbaceous plants.

## Acknowledgements

This project was supported by the University of Georgia, Georgia Research Alliance (C.R.B.) and Georgia Seed Development (C.R.B.), National Science Foundation MCB-2309665 (C.R.B and C.L.). T.T.T.D. receives funding from NSERC Alliance Collaboration (ALLRP 571673 – 21) and Catalyst (ALLRP 579871 – 22). T.T.T. Dang is grateful for the Michael Smith Health BC Scholar Award (SCH-2020-0401). Sequencing was performed at the University of Minnesota Genomics Center (UMGC) and UC Davis DNA Technologies and Expression Analysis Core. We thank Kathrine Mailloux for high molecular weight DNA isolation and Sophia Jones for plant care.

## Author contributions

CL, C.R.B., and T.T.T.D. designed the study. V.H.B performed metabolite profiling. C.L. generated single nuclei multiome libraries. L.H.C. performed cDNA sequencing. J.C.W. performed single cell library construction and quality control. B.V. performed genome assembly and data management. J.P.H. performed genome annotation. C.L. and V.H.B wrote the manuscript with input from all authors.

## Data Availability

All sequencing data associated with this study are available at the National Center for Biotechnology Institute Sequence Read Archive BioProject PRJNA1243444. Genome assembly, annotation, and Seurat objects for single cell multiome experiments are available via the digital repository figshare. Custom codes can be found at https://github.com/cxli233/Camptotheca_single_cell.

## Supporting Information

1) Supplemental methods. 2) Supplementary Figures S1-S5. 3) Supplementary Table S1-S9.

## Methods

### Plant growth conditions

One-year-old *C. acuminata* plants were bought from Planten Tuin Esveld, Netherlands. Plants were cultivated in a standard soil mix in a greenhouse at 20-25°C (daytime) and 17-20°C (nighttime) with 40-70% humidity under the minimum artificial light of 12 hrs at Jena Germany. At Athens, U.S.A., *C. acuminata* trees were grown in a greenhouse at 27-29°C (day time) and 17-20°C (night time) with 15 hrs light and natural humidity.

### *C*. *acuminata* metabolite profiling

Plant tissues (meristem, axillary bud, young leaf, mature leaf, young stem, mature stem, root) were collected and snap frozen in liquid nitrogen. Mortar and pestle were used to grind the tissues. Metabolites were extracted from the tissues using extraction solvent (85% methanol, 15% water, 0.1% formic acid) with the ratio of 100 mg tissues:1 ml extraction solvent. The extracts were filtered through 0.22 µm PTFE syringe filters. Extracts were diluted 100-fold with extraction buffer containing 50 nM catharanthine as an internal standard before LC-MS analysis.

### DNA isolation, sequencing, and genome assembly

High molecular weight DNA was isolated from immature leaf tissue using the Takara Nucleobond HMW DNA kit (Takara, Kusatsu, Shiga, Japan). Immature leaves were harvested from a tree two meters in height, grown in a greenhouse under the following conditions: 28ºC day, 19ºC night, with 15 hours of light at an intensity of 600 µM. Using the PacBio SMRTbell Template Prep Kit 3.0 (Menlo Park, CA), a single library was prepared and sequenced by the University of Minnesota Genomics Center. Sequencing was performed on a PacBio Revio instrument producing 47.4 Gb of HiFi data with a N50 of 18.5-kb, and an estimated coverage of 117x. PacBio HiFi reads were assembled using hifiasm v0.19.9-r616 (Cheng et al. 2024) with the homozygous coverage parameter set to 119. Contigs less than 50-kb were discarded using SeqKit (v2.8.2) (Shen et al. 2016) followed by two rounds of scaffolding using the scaffold tool in Rag-Tag (v2.1.0) (Alonge et al. 2019) with the previously published *C. acuminata* v3.0 assembly (Kang et al. 2021). Assembly metrics were calculated at each step using assembly-stats (v1.0.1) (github.com/sanger-pathogens/assembly-stats). PacBio HiFi reads and the genome assembly were checked for contamination using Kraken 2 (v2.1.3) (Wood, Lu, and Langmead 2019) with the PlusPFP database (k2_pluspfp_20240605, https://benlangmead.github.io/aws-indexes/k2).

BUSCO (v5.7.1) (Manni et al. 2021) was run at each step in the assembly process with the embryophyta_odb10 database to verify completeness of the genome assembly. Additional QC was performed by aligning the genome assembly to the *C. acuminata* v3.0 assembly (Kang et al. 2021) using minimap2 (v2.17-r941) (H. Li 2018) and the alignments visualized using D-Genies (Cabanettes and Klopp 2018).

### cDNA library construction, long reads sequencing, and processing

Mature leaf, shoot apex, stem, and root were used for RNA isolation using a modified hot borate method (Wan and Wilkins 1994). mRNA was isolated using the Dynabead mRNA Purification Kit (Thermo Fisher Scientific, 61011, Waltham, MA) and used as input into the Oxford Nanopore Technologies (ONT) SQK-PCB109 kit (ONT, Oxford, UK) to construct cDNA libraries which were then sequenced on R9 FLO-MIN106 Rev D flowcells on a MinION. Dorado (v0.7.3) github.com/nanoporetech/dorado) employing the SUP model (dna_r9.4.1_e8_sup@v3.6) to base-call the data with the following parameters also enabled; --no-trim --min-qscore 10 --no-classify.

### Genome Annotation

The genome assembly was repeat masked by first creating a custom repeat library (CRL) for the genome. Repeats were first identified with RepeatModeler (v2.03) (Flynn et al. 2020) and the version 3 genome assembly (Kang et al. 2021) and protein coding genes were filtered from the repeat database using ProtExcluder (v1.2) (Campbell et al. 2014) to create a CRL. The CRL was then combined with Viridiplantae repeats from RepBase (v20150807) (Bao, Kojima, and Kohany 2015) to generate the final CRL. The genome assembly was repeat-masked using the final CRL and RepeatMasker (v4.1.2-p1) (Tarailo□Graovac and Chen 2009) using the parameters -e ncbi -s-nolow -no_is -gff.

RNA-seq libraries were processed for genome annotation by first cleaning with Cutadapt (v2.10) (Martin 2011) using a minimum length of 50-nt and quality cutoff of 10 then aligning the cleaned reads to the genome using HISAT2 (2.1.0) (D. Kim et al. 2019). ONT cDNA reads were processed with Pychopper (v2.5.0) (github.com/epi2me-labs/pychopper) and trimmed reads greater than 500-nt were aligned to the genome using minimap2 (v2.17-r941) (H. Li 2018) with a maximum intron length of 5,000-nt. The aligned RNA-seq and ONT cDNA reads were each assembled using Stringtie (v2.2.1) (Kovaka et al. 2019) and transcripts less than 500-nt were removed.

Initial gene models were created using BRAKER2 (v2.1.6) (Hoff et al. 2019) using the softmasked genome assembly and the aligned RNA-seq libraries as hints. The gene models were then refined using two rounds of PASA2 (v2.5.2) (Haas 2003) to create a working gene model set. High-confidence gene models were identified from each working gene model by filtering out gene models without expression evidence, or a PFAM domain match, or were a partial gene model or contained an interior stop codon. Functional annotation was assigned by searching the working gene model proteins against the TAIR (v10) (Lamesch et al. 2012) database and the Swiss-Prot plant proteins (release 2015_08) database using BLASTP (v2.12.0) (Camacho et al. 2009) and the PFAM (v35.0) (El-Gebali et al. 2019) database using PfamScan (v1.6) and assigning the annotation based on the first significant hit. Visualization of synteny between v3 and v4 annotations were performed using GENESPACE (Lovell et al. 2022).

### Nuclei isolation

Nuclei isolation was performed as described previously (C. Li et al. 2022) with 0.0375% Triton-X-100 and no EDTA in the nuclei isolation buffer. Half a gram of immature leaves (∼3 cm in length) were chopped vigorously on ice on a petri dish in nuclei isolation buffer for exactly 2 min. The lysate was filtered through 100 µm and 40 µm sieves, which was then passed through a 20 µm sieve twice. Nuclei were stained with 4’,6-diamidino-2-phenylindole (DAPI) and sorted using a Moflo Astrios EQ flow cytometer at the UGA Cytometry Shared Resource Laboratory. At least 150,000 nuclei were sorted into 100 µL of nuclei buffer (part of the 10x Genomics Single Cell Multiome Kit). Nuclei were then centrifuged at 200 g for 5 min and resuspended in 50 µL nuclei buffer. The integrity of the nuclei was visually inspected using a fluorescence microscope. Single cell multi-ome libraries were constructed according to manufacturer’s recommendation (10x Genomics, Single Cell Multiome ATAC + Gene Expression Kit).

### Single nuclei RNA-seq processing

Single nuclei RNA-seq libraries were processed using Cutadapt (v3.5) (Martin 2011) with the following parameters: -q 30 -m 30 --trim-n -n 2 -g AAGCAGTGGTATCAACGCAGAGTACATGGG -a “A{20}”. The pairing of the reads was restored using SeqKit (v0.16.1) *pair* (Shen et al. 2016). Paired reads were aligned and quantified using STARsolo (Kaminow, Yunusov, and Dobin 2021), with the following parameters: -- alignIntronMax 5000 --soloUMIlen 12 --soloCellFilter EmptyDrops_CR --soloFeatures GeneFull --soloMultiMappers EM --soloType CB_UMI_Simple, and --soloCBwhitelist using the latest 10x Genomics whitelist of multiome barcodes. Gene-barcode matrices were analyzed with Seurat (v4) (Hao et al. 2021) for downstream analysis. Removal of low-quality nuclei and suspected multiplets was performed using the distributions of UMI counts and detected genes.

### Single nuclei RNA-seq analyses

Biological replicates were integrated using the ‘IntegrateData()‘ function in Seurat using the top 3,000 variable genes. Uniform manifold approximation and projection (UMAP) were performed after a principal component analysis (PCA) using the following parameters: dims = 1:30, min.dist = 0.001, repulsion.strength = 1, n.neighbors = 30, spread = 1. Clustering of cells was performed with a resolution of 0.5. Expression of previously studied biosynthetic genes were plotted across each cell cluster. For cell type classification, we used a manually curated marker gene list for mesophyll, epidermis, and vasculature, using previously established marker genes from Arabidopsis (J.-Y. Kim et al. 2021; Lopez-Anido et al. 2021). For dot-plot style expression heat maps, average expression of genes was calculated as the average Z-score of log-transformed normalized expression values across cell clusters and cell types. Dot sizes indicated the percentage of cells where a given gene is expressed (> 0 reads) in each cell type or cell cluster. *De novo* marker identification were performed using the ‘FindAllMarkers()‘ function in Seurat using the following options: only.pos = T, min.pct = 0.25, logfc.threshold = 0.25. Expression matrix (logCPM values) was generated at the cell cluster level, where each row is a gene, and each column is a cell cluster.

### Single nuclei ATAC-seq processing

Single nuclei ATAC-seq data were processed using the 10x Genomics Cell Ranger ARC pipeline (https://www.10xgenomics.com/software). The peak bed files for biological replicates were sorted and merged using BEDTools (v2.30) *merge* (Quinlan and Hall 2010). This common set of peaks was used to process both biological replicates. The ‘atac_fragments.tsv.gz’ files from the Cell Ranger ARC output were used for downstream analyses using Signac (v1.6.0) (Stuart et al. 2021) and Seurat (v4) (Hao et al. 2021). Nuclei were filtered for > 1000 peaks/nuclei, > 2000 fragments/nuclei, and fraction of fragments in peaks > 0.25. For data integration, the replicates were merged first, then integrated using the ‘IntegrateEmbeddings()‘ function in Signac using the “lsi” dimension reduction. Integration with the gene expression assay was performed by first filtering for shared nuclei in both gene expression and chromatin assays, after which the integrated ATAC-seq object was adjoined to the integrated RNA-seq object as a chromatin assay. By doing so, the cell cluster and cell type assignment information is transferred to the ATAC-seq assay. UMAP visualization for ATAC-seq was performed using the following parameters: reduction = “lsi”, dims = 2:30, min.dist = 0.001, repulsion.strength = 1, n.neighbors = 30, spread = 1. Joint UMAP visualization was done using the ‘FindMultiModalNeighbors()‘ functions in Signac. ATAC-seq coverage around genes and peaks was calculated and visualized using deepTools (v3.5.1) (Ramírez et al. 2014).

### Marker peak and motif overrepresentation analyses

Marker peaks for epidermis and idioblast were detected using the ‘FindMarkers()‘ function in Seurat after setting the default assay of the multiome object to chromatin accessibility, using the following parameters: only.pos = T, test.use = “LR”, min.pct = 0.05, latent.vars = ‘nCount_peaks’, group.by = “cell_type”. Only peaks with adjusted p-values < 0.05 were used for downstream analyses. Genes adjacent to STR+ marker peaks were defined using BEDTools (v.2.30) *closest*, with -d option. Only genes within 2-kb to a STR+ marker peak were used in visualization. DNA sequence of STR+ maker peaks were extracted using BEDTools (v.2.30) *getfasta* and subjected to *de novo* motif discovery using MEME (v5.4.1) (Bailey et al. 2009): using the following parameters: -dna -revcomp -mod anr -nmotifs 10 -minw 5 -maxw 12 -evt 0.01. Similar motifs were identified from JASPAR2020 collection (Fornes et al. 2019).

## References

Fornes, Oriol, Jaime A Castro-Mondragon, Aziz Khan, Robin van der Lee, Xi Zhang, Phillip A Richmond, Bhavi P Modi, et al. 2019. “JASPAR 2020: Update of the Open-Access Database of Transcription Factor Binding Profiles.” Nucleic Acids Research, November, gkz1001. 10.1093/nar/gkz1001.

Kim, Ji-Yun, Efthymia Symeonidi, Tin Yau Pang, Tom Denyer, Diana Weidauer, Margaret Bezrutczyk, Manuel Miras, et al. 2021. “Distinct Identities of Leaf Phloem Cells Revealed by Single Cell Transcriptomics.” The Plant Cell 33 (3): 511–30. 10.1093/plcell/koaa060.

Li, Chenxin, Joshua C. Wood, Anh Hai Vu, John P. Hamilton, Carlos Eduardo Rodriguez Lopez, Richard M. E. Payne, Delia Ayled Serna Guerrero, et al. 2023. “Single-Cell Multi-Omics in the Medicinal Plant Catharanthus Roseus.” Nature Chemical Biology 19 (8): 1031–41. 10.1038/s41589-023-01327-0.

Lorence, Argelia, and Craig L. Nessler. 2004. “Camptothecin, over Four Decades of Surprising Findings.” Phytochemistry 65 (20): 2735–49. 10.1016/j.phytochem.2004.09.001.

Nguyen, Tuan-Anh M., Trinh-Don Nguyen, Yuen Yee Leung, Matthew McConnachie, Oleg Sannikov, Zhicheng Xia, and Thu-Thuy T. Dang. 2021. “Discovering and Harnessing Oxidative Enzymes for Chemoenzymatic Synthesis and Diversification of Anticancer Camptothecin Analogues.” Communications Chemistry 4 (1): 177. 10.1038/s42004-021-00602-2.

Sadre, Radin, Maria Magallanes-Lundback, Sujana Pradhan, Vonny Salim, Alex Mesberg, A. Daniel Jones, and Dean DellaPenna. 2016. “Metabolite Diversity in Alkaloid Biosynthesis: A Multilane (Diastereomer) Highway for Camptothecin Synthesis in Camptotheca Acuminata.” The Plant Cell 28 (8): 1926–44.

Salim, Vonny, A Daniel Jones, and Dean DellaPenna. 2018. “Camptotheca Acuminata 10-Hydroxycamptothecin O-Methyltransferase: An Alkaloid Biosynthetic Enzyme Co-Opted from Flavonoid Metabolism.” The Plant Journal 95 (1): 112–25.

## Reference

Alonge, Michael, Sebastian Soyk, Srividya Ramakrishnan, Xingang Wang, Sara Goodwin, Fritz J. Sedlazeck, Zachary B. Lippman, and Michael C. Schatz. 2019. “RaGOO: Fast and Accurate Reference-Guided Scaffolding of Draft Genomes.” Genome Biology 20 (1): 224. 10.1186/s13059-019-1829-6.

Bailey, T. L., M. Boden, F. A. Buske, M. Frith, C. E. Grant, L. Clementi, J. Ren, W. W. Li, and W. S. Noble. 2009. “MEME SUITE: Tools for Motif Discovery and Searching.” Nucleic Acids Research 37 (Web Server): W202–8. 10.1093/nar/gkp335.

Bao, Weidong, Kenji K. Kojima, and Oleksiy Kohany. 2015. “Repbase Update, a Database of Repetitive Elements in Eukaryotic Genomes.” Mobile DNA 6 (1): 11. 10.1186/s13100-015-0041-9.

Cabanettes, Floréal, and Christophe Klopp. 2018. “D-GENIES: Dot Plot Large Genomes in an Interactive, Efficient and Simple Way.” Edited by Thomas Tullius. PeerJ 6 (June):e4958. 10.7717/peerj.4958.

Camacho, Christiam, George Coulouris, Vahram Avagyan, Ning Ma, Jason Papadopoulos, Kevin Bealer, and Thomas L Madden. 2009. “BLAST+: Architecture and Applications.” BMC Bioinformatics 10 (1): 421. 10.1186/1471-2105-10-421.

Campbell, Michael S., MeiYee Law, Carson Holt, Joshua C. Stein, Gaurav D. Moghe, David E. Hufnagel, Jikai Lei, et al. 2014. “MAKER-P: A Tool Kit for the Rapid Creation, Management, and Quality Control of Plant Genome Annotations.” Plant Physiology 164 (2): 513–24. 10.1104/pp.113.230144.

Cheng, Haoyu, Mobin Asri, Julian Lucas, Sergey Koren, and Heng Li. 2024. “Scalable Telomere-to-Telomere Assembly for Diploid and Polyploid Genomes with Double Graph.” Nature Methods 21 (6): 967–70. 10.1038/s41592-024-02269-8.

El-Gebali, Sara, Jaina Mistry, Alex Bateman, Sean R Eddy, Aurélien Luciani, Simon C Potter, Matloob Qureshi, et al. 2019. “The Pfam Protein Families Database in 2019.” Nucleic Acids Research 47 (D1): D427–32. 10.1093/nar/gky995.

Flynn, Jullien M., Robert Hubley, Clément Goubert, Jeb Rosen, Andrew G. Clark, Cédric Feschotte, and Arian F. Smit. 2020. “RepeatModeler2 for Automated Genomic Discovery of Transposable Element Families.” Proceedings of the National Academy of Sciences 117 (17): 9451–57. 10.1073/pnas.1921046117.

Haas, B. J. 2003. “Improving the Arabidopsis Genome Annotation Using Maximal Transcript Alignment Assemblies.” Nucleic Acids Research 31 (19): 5654–66. 10.1093/nar/gkg770.

Hao, Yuhan, Stephanie Hao, Erica Andersen-Nissen, William M. Mauck, Shiwei Zheng, Andrew Butler, Maddie J. Lee, et al. 2021. “Integrated Analysis of Multimodal Single-Cell Data.” Cell 184 (13): 3573-3587.e29. 10.1016/j.cell.2021.04.048.

Hoff, Katharina J, Alexandre Lomsadze, Mark Borodovsky, and Mario Stanke. 2019. “Whole-Genome Annotation with BRAKER.” Gene Prediction: Methods and Protocols, 65–95.

Kaminow, Benjamin, Dinar Yunusov, and Alexander Dobin. 2021. “STARsolo: Accurate, Fast and Versatile Mapping/Quantification of Single-Cell and Single-Nucleus RNA-Seq Data.” bioRxiv, May. 10.1101/2021.05.05.442755.

Kang, Minghui, Rao Fu, Pingyu Zhang, Shangling Lou, Xuchen Yang, Yang Chen, Tao Ma, Yang Zhang, Zhenxiang Xi, and Jianquan Liu. 2021. “A Chromosome-Level Camptotheca Acuminata Genome Assembly Provides Insights into the Evolutionary Origin of Camptothecin Biosynthesis.” Nature Communications 12 (1): 3531. 10.1038/s41467-021-23872-9.

Kim, Daehwan, Joseph M. Paggi, Chanhee Park, Christopher Bennett, and Steven L. Salzberg. 2019. “Graph-Based Genome Alignment and Genotyping with HISAT2 and HISAT-Genotype.” Nature Biotechnology 37 (8): 907–15. 10.1038/s41587-019-0201-4.

Kovaka, Sam, Aleksey V. Zimin, Geo M. Pertea, Roham Razaghi, Steven L. Salzberg, and Mihaela Pertea. 2019. “Transcriptome Assembly from Long-Read RNA-Seq Alignments with StringTie2.” Genome Biology 20 (1): 278. 10.1186/s13059-019-1910-1.

Lamesch, Philippe, Tanya Z. Berardini, Donghui Li, David Swarbreck, Christopher Wilks, Rajkumar Sasidharan, Robert Muller, et al. 2012. “The Arabidopsis Information Resource (TAIR): Improved Gene Annotation and New Tools.” Nucleic Acids Research 40 (D1): D1202–10. 10.1093/nar/gkr1090.

Li, Chenxin, Joshua C. Wood, Natalie C. Deans, Anne Frances Jarrell, Dionne Martin, Kathrine Mailloux, Yi-Wen Wang, and C. Robin Buell. 2022. “Nuclei Isolation Protocol from Diverse Angiosperm Species.” bioRxiv, November. 10.1101/2022.11.03.515090.

Li, Heng. 2018. “Minimap2: Pairwise Alignment for Nucleotide Sequences.” Edited by Inanc Birol. Bioinformatics 34 (18): 3094–3100. 10.1093/bioinformatics/bty191.

Lopez-Anido, Camila B., Anne Vatén, Nicole K. Smoot, Nidhi Sharma, Victoria Guo, Yan Gong, M. Ximena Anleu Gil, Annika K. Weimer, and Dominique C. Bergmann. 2021. “SingleCell Resolution of Lineage Trajectories in the Arabidopsis Stomatal Lineage and Developing Leaf.” Developmental Cell 56 (7): 1043-1055.e4. 10.1016/j.devcel.2021.03.014.

Lovell, John T, Avinash Sreedasyam, M Eric Schranz, Melissa Wilson, Joseph W Carlson, Alex Harkess, David Emms, David M Goodstein, and Jeremy Schmutz. 2022. “GENESPACE Tracks Regions of Interest and Gene Copy Number Variation across Multiple Genomes.” eLife 11 (September):e78526. 10.7554/eLife.78526.

Manni, Mosè, Matthew R Berkeley, Mathieu Seppey, Felipe A Simão, and Evgeny M Zdobnov. 2021. “BUSCO Update: Novel and Streamlined Workflows along with Broader and Deeper Phylogenetic Coverage for Scoring of Eukaryotic, Prokaryotic, and Viral Genomes.” Molecular Biology and Evolution 38 (10): 4647–54. 10.1093/molbev/msab199.

Martin, Marcel. 2011. “Cutadapt Removes Adapter Sequences from High-Throughput Sequencing Reads.” EMBnet 17 (1): 3.

Quinlan, Aaron R., and Ira M. Hall. 2010. “BEDTools: A Flexible Suite of Utilities for Comparing Genomic Features.” Bioinformatics 26 (6): 841–42. 10.1093/bioinformatics/btq033.

Ramírez, Fidel, Friederike Dündar, Sarah Diehl, Björn A. Grüning, and Thomas Manke. 2014. “deepTools: A Flexible Platform for Exploring Deep-Sequencing Data.” Nucleic Acids Research 42 (W1): W187–91. 10.1093/nar/gku365.

Shen, Wei, Shuai Le, Yan Li, and Fuquan Hu. 2016. “SeqKit: A Cross-Platform and Ultrafast Toolkit for FASTA/Q File Manipulation.” Edited by Quan Zou. PLOS ONE 11 (10): e0163962. 10.1371/journal.pone.0163962.

Stuart, Tim, Avi Srivastava, Shaista Madad, Caleb A. Lareau, and Rahul Satija. 2021. “Single-Cell Chromatin State Analysis with Signac.” Nature Methods 18 (11): 1333–41. 10.1038/s41592-021-01282-5.

Tarailo□Graovac, Maja, and Nansheng Chen. 2009. “Using RepeatMasker to Identify Repetitive Elements in Genomic Sequences.” Current Protocols in Bioinformatics 25 (1). 10.1002/0471250953.bi0410s25.

Wan, C. Y., and T. A. Wilkins. 1994. “A Modified Hot Borate Method Significantly Enhances the Yield of High-Quality RNA from Cotton (Gossypium Hirsutum L.).” Analytical Biochemistry 223 (1): 7–12. 10.1006/abio.1994.1538.

Wood, Derrick E., Jennifer Lu, and Ben Langmead. 2019. “Improved Metagenomic Analysis with Kraken 2.” Genome Biology 20 (1): 257. 10.1186/s13059-019-1891-0.

