## Supplementary figures for "Single cell multi-omics reveals rare biosynthetic cell types in the medicinal tree *Camptotheca acuminata*"

**Supplementary Figures and Legend**

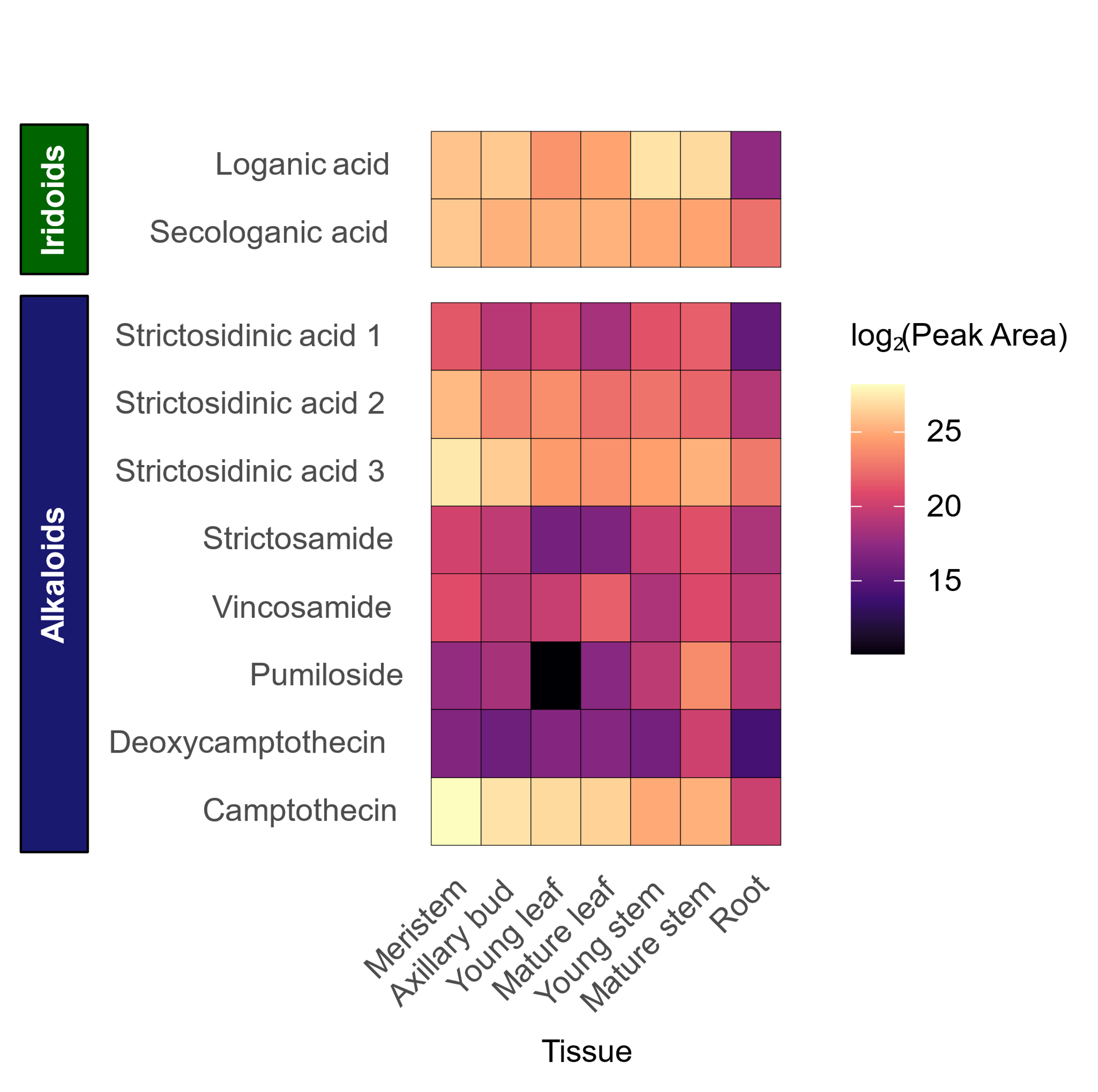

**Fig. S1. Key monoterpene indole alkaloids (MIA) of *Camptotheca acuminata.***

Relative abundances of key iridoids and MIAs across major organs.

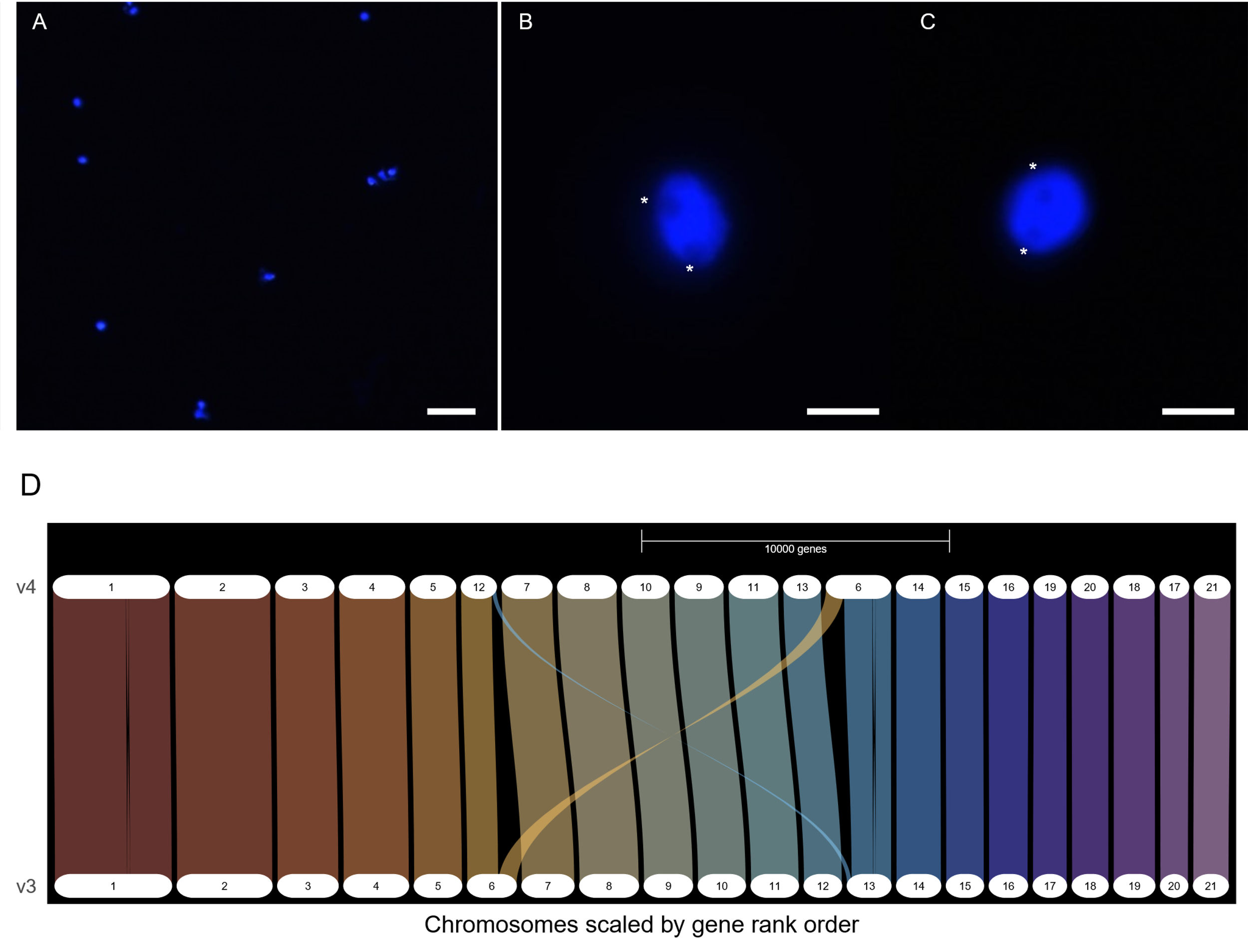

**Fig. S2. *Camptotheca acuminata* v4 genome assembly and annotation and nuclei preparation for single nuclei omics.**

(A). Isolated nuclei from immature leaves (20× objective lens). Bar: 40 µm.

(B)-(C). Isolated nuclei from immature leaves (100× objective lens). Bar: 10 µm. Asterisks indicate nucleoli.

(D). Riparian plots comparing v4 and v3 (Kang et al. 2021) genome assembly and annotation generated using GENESPACE (Lovell et al. 2022). Chromosomes are scaled by gene order, not by physical size.

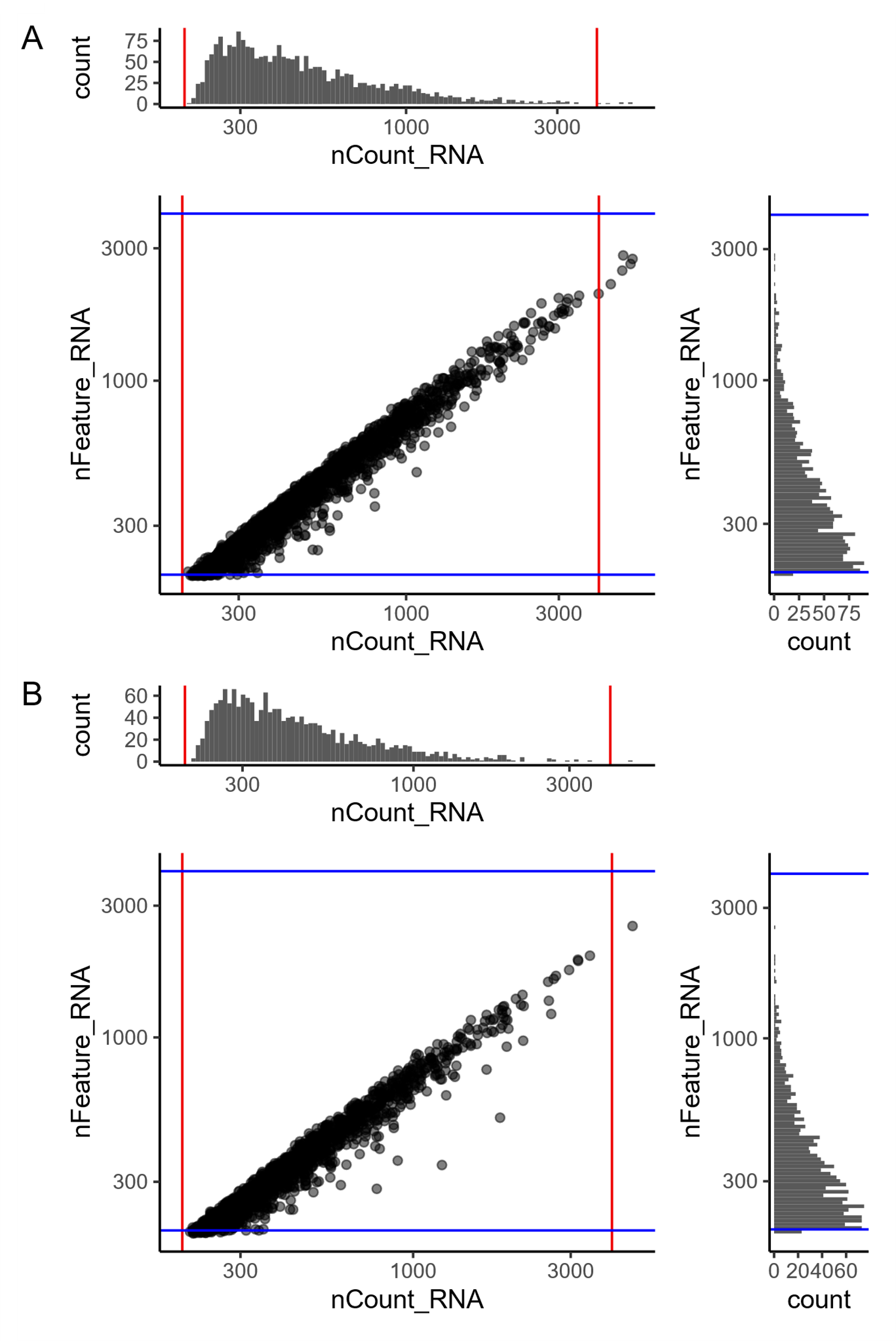

**Fig. S3. Characterization of leaf single nuclei RNA-seq libraries.**

(A)-(B). Scatter plots of nCount_RNA (UMI count) and nFeature_RNA (number of detected genes) for two biological replicates from immature leaf samples. Both axes are in log_10_ scale. The distribution of UMI count and gene count are shown above and to the right of the scatter plots, respectively. Only nuclei between the red and blue lines are used for downstream analyses.

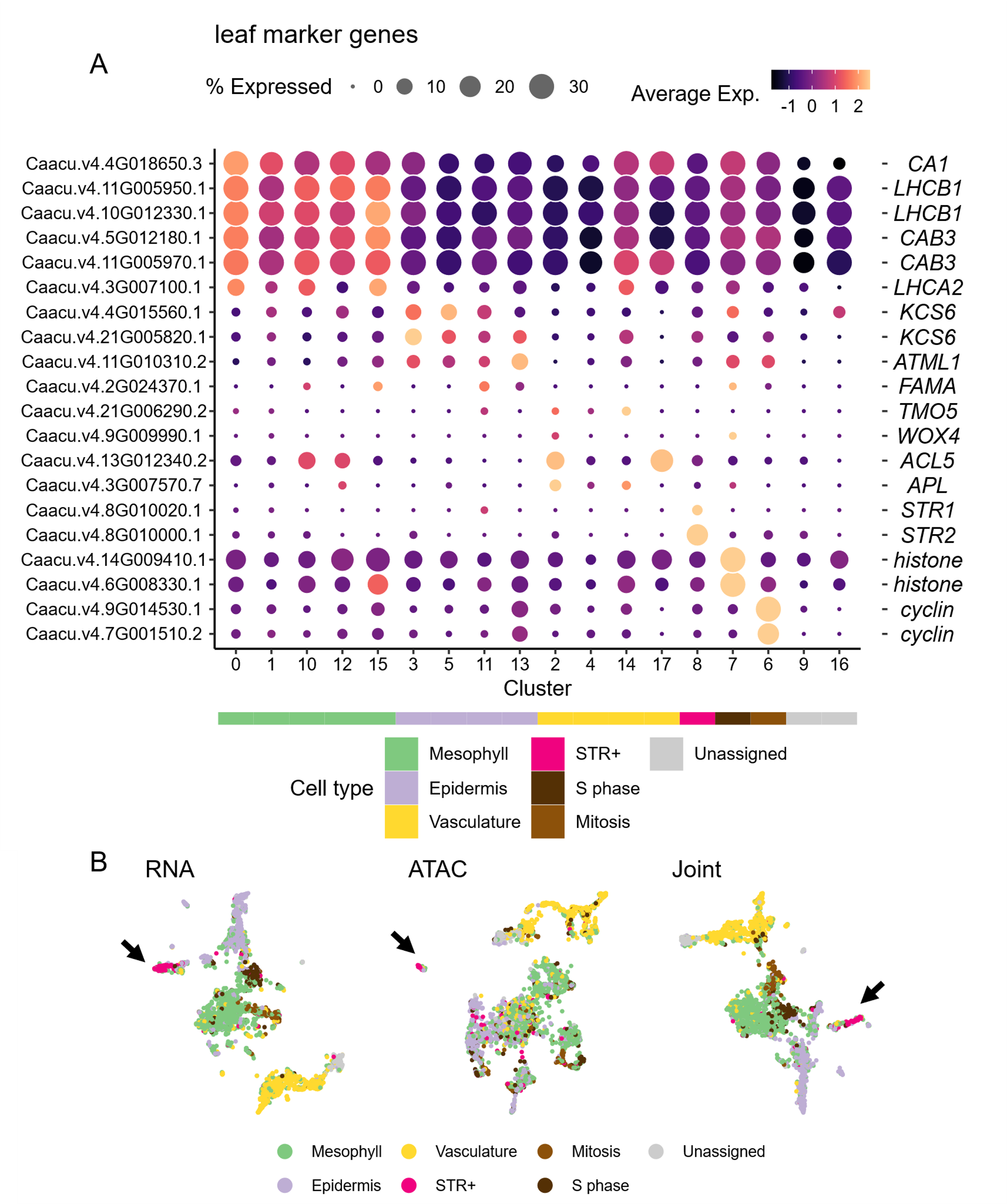

**Fig. S4. Maker genes and multimodal integration for leaf single nuclei omics datasets.**

(A). Gene expression heatmap of marker genes across leaf cell clusters. Color scale shows the average scaled expression of each gene at each cell cluster. Dot size indicates the percentage of cells where a given gene is detected. The predicted cell type for each cell cluster is annotated by the color strip below the x-axis. Gene symbols are shown on the right.

(B). Uniform Manifold Approximation and Projection (UMAP) of nuclei containing both high-quality RNA-seq and ATAC-seq data color coded by cell types, with the same color palette as in (A). From left to right: UMAP based on gene expression assay, chromatin accessibility assay, and joint analysis. Arrows indicate STR+ cells.

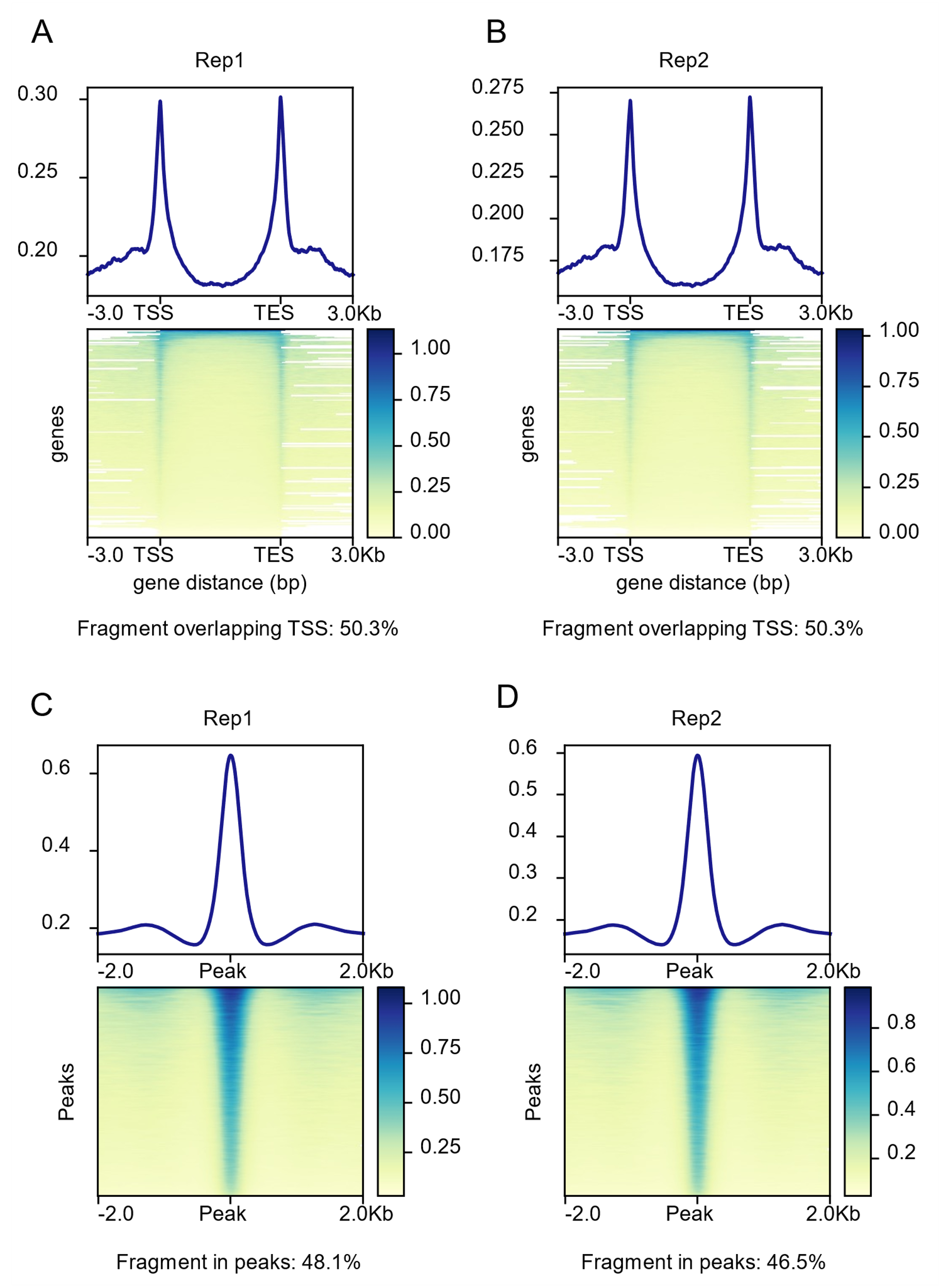

**Fig. S5. Characterization of leaf single nuclei ATAC-seq libraries.**

(A)-(B). Metagene plot (top) and heatmap (bottom) showing coverage of ATAC-seq fragments around genes for each biological replicate. In the heatmap, each row is a gene. TSS. Transcription start site. TES. Transcription end site. The percentage of fragments overlapping TSSs are listed below the heatmap.

(C)-(D). Metagene plot (top) and heatmap (bottom) showing coverage of ATAC-seq fragments around ATAC-seq peaks for each biological replicate. In the heatmap, each row is an ATAC-seq peak. The percentage of fragments overlapping peaks are listed below the heatmap.

References

Kang, Minghui, Rao Fu, Pingyu Zhang, Shangling Lou, Xuchen Yang, Yang Chen, Tao Ma, Yang Zhang, Zhenxiang Xi, and Jianquan Liu. 2021. “A Chromosome-Level Camptotheca Acuminata Genome Assembly Provides Insights into the Evolutionary Origin of Camptothecin Biosynthesis.” *Nature Communications* 12 (1): 3531. https://doi.org/10.1038/s41467-021-23872-9.

Lovell, John T, Avinash Sreedasyam, M Eric Schranz, Melissa Wilson, Joseph W Carlson, Alex Harkess, David Emms, David M Goodstein, and Jeremy Schmutz. 2022. “GENESPACE Tracks Regions of Interest and Gene Copy Number Variation across Multiple Genomes.” *eLife* 11 (September):e78526. https://doi.org/10.7554/eLife.78526.
